# Demographic and genetic impacts of powdery mildew in a young oak cohort

**DOI:** 10.1101/2023.06.22.546164

**Authors:** Benoit Barrès, Cyril Dutech, Gilles Saint-Jean, Catherine Bodénès, Christian Burban, Virgil Fiévet, Camille Lepoittevin, Pauline Garnier-Géré, Marie-Laure Desprez-Loustau

## Abstract

The demographic and genetic impacts of powdery mildew on the early stages of an oak population were studied in an *ad hoc* field design with two disease exposures. This enabled a detailed phenotypic monitoring of 1,733 emerging individuals from 15 progenies over nine years, and the genotyping of 68% of them. The pathogen induced high levels of seedling mortality several years after sowing, associated with reduced growth and capacity to overwinter. The probability of juvenile survival could be predicted from mean disease severity in early years and acorn weight. Fast-growing families showed the highest survival rate under both natural and protected disease exposure. Correlatively, no equalizing effect of increased powdery mildew pressure on the relative contribution of mother trees to the next generation could be detected. Contrary to a possible trade-off hypothesis between growth and defense, family height potential was not negatively related to disease resistance across the studied oak mother trees. Overall, our results suggest that in *Quercus robur* natural populations, infection levels (related to resistance *sensu stricto*) may be less determinant than growth-related tolerance to infection for the fate of seedlings. However, an equalizing effect of powdery mildew on relative oak genotype performances cannot be excluded at later stages since such an effect was already visible on height. Average genomic diversity was not significantly affected by mortality associated with powdery mildew. However, our study brings support to a deleterious effect of very low individual heterozygosity on the probability of survival across the different families. Finally, our study points to a few candidate genes for several fitness-related traits.

## Introduction

Seedling establishment and early growth stages are crucial phases in the tree life cycle. Most forest tree species show a typical concave mortality curve, characterized by a very high juvenile mortality (*e.g.*, Harcombe 1987; Peñuelas et al 2007; Petit & Hampe 2005; Kelly 2002). Under natural conditions, both abiotic and biotic factors affect tree seedling survival, in addition to stochastic processes (*e.g.*, Shibata et al 2010; Petritan et al 2014; Martini et al 2019). Among biotic factors, many pathogens may affect seedling, and more generally juvenile survival. Seedlings and saplings are especially susceptible to pathogens due to their mostly non-woody tissues, both in roots and stems (Dominguez-Begines et al 2020; Jankowiak et al 2022). For example, Augspurger (1984) reported that damping-off pathogens (soilborne fungi and oomycetes) accounted for the largest proportion of seedling deaths within the first year in several species of tropical forest trees.

By their negative effect on the individual fitness of their host (by definition), pathogens can strongly affect plant population demographic patterns. On the other hand, at community level, their positive role in maintaining between and within species diversity has received increasing support (Dobson & Crawley 1994; Alexander 2010; Mordecai 2011; Bever et al 2015). The impact of pests and pathogens on seedlings has been extensively studied as a possible mechanism promoting tree species coexistence (maintenance of spatial diversity) in species-rich tropical forests. According to the Janzen-Connell model, species-specific herbivores and pathogens provide a frequency-dependent spacing (thus diversifying) mechanism by causing increased mortality of seedlings growing at a short distance from their mother tree (Janzen 1970; Connell 1971; Summers et al 2003). Many studies have provided support to this model both in tropical and temperate environments (Packer & Clay 2000; Bell et al 2006; Yamazaki et al 2008; Terborgh 2020), although the magnitude and generality of Janzen-Connell effects are still a matter of debate (Song et al 2021). Such frequency-dependent and density-dependent processes are especially important for specialized pathogens, as in Janzen-Connell effects, or in co-evolutionary dynamics at population level (Mundt et al 2008; Parker & Gilbert 2018; Burdon & Laine 2019). Pathogens may also affect competitive interactions between genotypes, between or within species, in a non-frequency dependent or density-dependent manner, by causing a differential cost on the fitness of the competing plants (Mundt et al 2008; Creissen et al 2016). For example, foliar diseases have a debilitating effect on highly infected seedlings, which may result in a competitive disadvantage in presence of less affected neighbors (Wiener 1990; Gilbert 2002; Power & Mitchell 2004). When the competitively dominant genotypes in the absence of disease experience a greater cost to disease than less competitive genotypes in the presence of pathogens, pathogens reduce fitness differences and therefore promote plant diversity (Mordecai 2011). This occurs when the fast growing/strongest competitors are the most vulnerable to pathogens (Summers et al 2003; Bever et al 2015; Cope et al 2021). The prevailing hypothesis in the literature to explain this negative correlation is the growth-defense trade-off concept, based on the premise that defense is costly thus requires allocation of resources at the expense of growth (Monson et al 2022). Growth-defense trade-offs have been reported at inter-and intra-specific level in many groups of plants under various environments, including for tree species (Lind et al 2013; Heckman et al 2019; Kruger et al 2020; Cope et al 2021).

Studies on the impact of pathogens on plant populations have been extensively performed in an agricultural context, in relation to yield losses (*e.g.*, Savary et al 2019). Studies in natural systems are fewer, and mainly focused on some model systems (Burdon & Thrall 2014), *e.g.*, flax rust (Thrall et al 2012), Arabidopsis pathogens (Creissen et al 2016), anther smut of Silene (Bernasconi et al 2009), Plantago powdery mildew (Laine 2004; Safdari et al 2021). In this study, we aimed to characterize the impacts of powdery mildew on fitness-related traits and genetic diversity during the early life-stages of an oak cohort. Powdery mildew is one of the most important diseases on temperate oaks in Europe, in particular pedunculate oak, *Quercus robur* (Mougou et al 2008; Lonsdale 2015). In Europe, powdery mildew was shown to be associated with a complex of cryptic (morphologically similar) species, of which *Erysiphe alphitoides* (Griffon & Maubl.) U. Braun & S. Takam is nowadays the most prevalent throughout Europe, often in mixture with *Erysiphe quercicola* in southern Europe S. Takam. & U. Braun and with *Erysiphe hypophylla* (Nevod.) U. Braun & Cunningt. in northern Europe (Mougou et al 2008; Desprez-Loustau et al 2018; Gross et al 2021). Demeter et al (2021) suggested that powdery mildew could be one of the major factors involved in regeneration failures in pedunculate oak throughout Europe. Seedlings and young trees, with a relatively high amount of young, succulent, fast growing tissues, are especially susceptible to disease (Pap et al 2012; Marçais & Desprez-Loustau 2014). A significant negative effect of powdery mildew on height and radial growth of oak saplings was demonstrated in comparison with controls protected by fungicide applications (Pap et al 2012; Desprez-Loustau et al 2014). Powdery mildew, as an obligate parasite, derives nutrients produced by plant photosynthesis to its own benefit thanks to specialized feeding structures (called haustoria) that penetrate into living cells of the leaf parenchyma (Hewitt & Ayres 1976). As a consequence, several types of damage have been described: reduced net assimilation rate, reduced height and radial growth, greater susceptibility to frost (Hajji et al 2009; Marçais & Desprez-Loustau 2014; Pap et al 2014; Bert et al 2016). However, how the impacts of powdery mildew scale up at oak population level have rarely been explored.

The spatial, demographic, and genetic structure of oak populations (especially *Q. robur* and *Q. petraea*) has nevertheless received much attention owing to the ecologic, cultural and economic importance of these species in Europe (*e.g.*, Kremer & Petit 1993; Streiff et al 1998; Gömöry et al 2001; Vakkari et al 2006; Kesić et al 2021). Overall, oak populations exhibit a high level of genetic diversity, with no significant or little differences among cohorts of different ages in the same stand (Vranckx et al 2014a from adults to established seedlings; Gerzabek et al 2020 from emergence to 3-year old seedlings). In a natural context, the various biotic and abiotic factors affecting oak seedling recruitment can vary in space and time (*e.g.*, Crawley & long 1995; Alberto et al 2011; Gerzabek et al 2020). The diversity and fluctuation of selective pressures acting on different genetic components have been proposed as possible explanations for the maintenance of genetic diversity in plant populations (Ennos 1983; Delph & Kelly 2014).

Genetic changes in plant populations under pathogen pressure have been reported in a few pathosystems (Thrall et al 2012). In this case, with alleles being selected due to their positive association with a greater resistance and/or tolerance to the disease, it may be possible to identify some of these variants using an association genetics approach. Genome Wide Association Studies (GWAS) are a powerful tool to link phenotypic variation with genetic polymorphisms, allowing the identification of the underlying biological mechanisms (Korte & Farlow 2013; Tibbs Cortes et al 2021). High quality genomic resources are now available for *Q. robur* (Lepoittevin et al 2015; Plomion et al 2018; Lang et al 2021). Both genetic variation among families (Desprez-Loustau et al 2014) and putative candidate genomic regions for oak susceptibility to powdery mildew (Bartholomé et al 2020) were previously demonstrated in independent studies. Together with a very high genomic diversity within oak populations and a rapid decay of linkage disequilibrium among variants across the oak genome (Lang et al 2021), these species characteristics provide an advantageous setting for performing GWAS.

Our general objective was to characterize the demographic and genetic impact of powdery mildew in the early stages of an oak population. We used an original experimental field design with two levels of powdery mildew exposure and a half-sib family genetic structure, that we analyzed with a large range of methods, in order to address the following questions:

1. How does powdery mildew affect juvenile survival? Phenotypic monitoring was carried out during the first nine years after sowing. We analyzed the effect of powdery mildew on the probability of seedling survival with various logistic regression models, and used structural equation modelling (SEM) to describe the multiple relationships between the measured phenotypic variables and survival.
2. Does the impact of powdery mildew, in terms of survival, vary among oak families, *i.e.*, does powdery mildew differentially affect the reproductive success of different oak mother trees? In particular, do the families performing best (*i.e.*, with greatest survival and growth) in conditions of low powdery mildew pressure also perform best in conditions of high powdery mildew pressure? We hypothesized that seedling and juvenile survival is strongly affected by early growth, this trait itself being sensitive to both maternal effects such as those due to acorn weight and average family or individual genotypic effects, but that growth could be negatively correlated with resistance to the pathogen (*i.e.*, associated with a growth-resistance trade-off).
3. As a consequence, does powdery mildew reduce fitness differences of mother trees, measured by the mean survival of their progenies, *i.e.*, has powdery mildew an equalizing effect? If so, is the surviving population more or less diverse in terms of family composition under high powdery mildew pressure than under low disease pressure?
4. Does powdery mildew impact the genetic diversity of the oak population, not only in terms of family composition? In order to address this question, a large number of emerging seedlings were genotyped at several hundred SNPs and heterozygosity statistics were calculated. The genetic diversity was then compared in the surviving and initial populations under the two contrasted disease pressures. Furthermore, we tested whether possible genetic changes could be associated with a difference in individual heterozygosity between dead and living seedlings, in agreement with the HFC (Heterozygosity-Fitness-Correlations) hypothesis (Vranckx et al 2014b).
5. Finally, given our experimental setting with a known family structure, can we detect significant genetic associations between some loci and seedling survival or other related traits (growth, infection)?

## Methods

### Experimental design

The experimental design and trait definitions were thoroughly described in the phenotypic monitoring study during the first three years (Desprez-Loustau et al 2014). Briefly, the progeny of 15 isolated (*i.e.* with no overlapping canopies) oak trees (*Q. robur*) was collected in 2008 in Cestas, France. *Q robur* is a diploid, monoecious, wind-pollinated tree species, with a highly outcrossing breeding system; it is a light-demanding species (especially at juvenile stages) with a moderate tolerance to drought, compared with other oaks such as *Q. petraea* (Eaton et al 2016). We thus considered a population made of 15 open-pollinated half-sib families. The weight of each acorn was recorded for its importance on the initial seedling developmental stage (Sánchez-Montes de Oca et al 2018). Acorns were then sown in April 2009 on a 10 x 10 cm grid in a field design with 9 unit plots, each containing 296 acorns, and distributed in 3 blocks (Supplementary material Figures S1 and S2). The field design was located in the INRAE experimental domain in Pierroton, Cestas, France. The unit plots were randomly attributed to one of two powdery mildew exposures: either Natural or Protected, *i.e.*, with a protection provided by myclobutanil (Dow AgroSciences, Sophia Antipolis, France), a fungicide authorized for usage in nurseries. There were six unit plots with natural exposure (corresponding to the pooled “Medium” and “High” disease treatments described in Desprez-Loustau et al 2014, which did not differ much in seedling infection and mortality) and three with fungicide (myclobutanil, a systemic fungicide which inhibits the ergosterol biosynthesis of the fungus) application. Although fungicide application limits the level of infection, it does not completely prevent the disease on treated trees. Acorns from different families were randomly distributed among plots, with 173 acorns per family on average (minimum=118; maximum=285; Supplementary material Figure S2).

At the end of each growing season from 2009 to 2012, survival was noted and height was measured for each individual. In following years, survival and height were assessed in early spring, a few weeks after budburst. Tree height was defined as the height of the highest living bud (*i.e.*, with leaves). Apical bud mortality occurred in some years, *i.e.*, the upper stem and branches did not show bud burst, resulting in a negative net annual height growth (Desprez-Loustau et al 2014).

Powdery mildew infection was monitored and scored on several occasions each year, particularly in the first years. To characterize annual disease severity, we used data corresponding to the highest annual infection score (generally from the last assessment at the end of the season, in September or October, but sometimes earlier, depending on the year’s powdery mildew epidemic dynamic). For example, in 2011, early and severe epidemics occurred, resulting in premature defoliation. Disease monitoring was therefore stopped in July. Disease severity assessments are therefore not directly comparable between years. No assessment was done in 2014 and 2015, during which infection was low. Powdery mildew infection was estimated visually by trained observers as a percentage of total infected leaf area for each individual. From the molecular analysis of various samplings in the experimental field, in different years (206 analyses in total), it was confirmed that the most prevalent species was *E. alphitoides* (more than 90% samples). *E. quercicola* was the other species detected, albeit at a much lower prevalence.

A late frost occurred in spring 2013, resulting in leaf damage in some seedlings. The occurrence of such damage was recorded as present/absent for each seedling. The details of the variables used in the statistical analyses are described in the Supplementary material, Table S1.

### Statistical analyses

#### Logistic and structural equation modelling for analysis at the individual level

In order to explore the impacts of powdery mildew on survival, we used two statistical approaches. First, logistic models were used to test the effects of different variables (see Supplementary material,

Table S2) on tree survival (“Survival (2017)”, dead or living). In the first and simplest model, two explanatory variables of survival were included: “Acorn weight” and “Powdery mildew exposure” (Natural exposure or Protected by fungicide, Model 1). In order to further detail the powdery mildew effects, we replaced the exposure variable by a quantitative variable corresponding to the mean disease severity over the first five years (“Mean infection (2009-2013)”, Model 2). Another model was also run with seedling height at the end of the first growing season in place of acorn weight (“Height in 2009”, Model 3). The binary (*i.e.*, yes/no) variable “Frost damage (2013)” was then added to Model 1 (Model 4). Finally, a full logistic model of survival included previously studied factors (powdery mildew exposure, acorn weight and frost damage), with a family effect and an interaction effect between powdery mildew exposure and family (Model 5).

Second, we used Structural Equation Modelling (SEM) in order to estimate multiple and interrelated dependencies among measured phenotypic variables and survival. Based on a pre-defined causal model, the SEM method gives quantitative estimates of direct and indirect effects of several inter-correlated variables on a variable of interest, according to different “paths”. Standardized coefficients (*i.e.*, relationships expressed in terms of standard deviations) are produced by the analysis, enabling the comparison of the relative strengths of the effects of different explanatory variables, along the different paths. In addition, the total effect of each explanatory variable on the variable of interest is broken down into its direct and indirect effects, according to the specified paths. In our case, the main objective was to understand how powdery mildew affects the final survival of seedlings (in 2017). The phenotypic variables used in SEM were the mean seedling infection over the first five growing seasons (2009-2013), the height at the end of 2012 and the presence of frost damage in early 2013. These variables were selected because mortality only started in 2014 thus phenotypic data needed for the model were available for almost all seedlings. We assumed that differences in infection and height in the first five years, as well as frost damage, were important determinants of subsequent survival. Moreover, susceptibility and growth expressed during the first five years are likely correlated with susceptibility and growth in subsequent years. Taking into account previous knowledge on powdery mildew, we constructed and tested a model with three “paths” (corresponding to potentially different mechanisms) relating disease severity (“Mean infection (2009-2013)”) to survival (“Survival 2017”, Figure 1).

**Figure 1:**
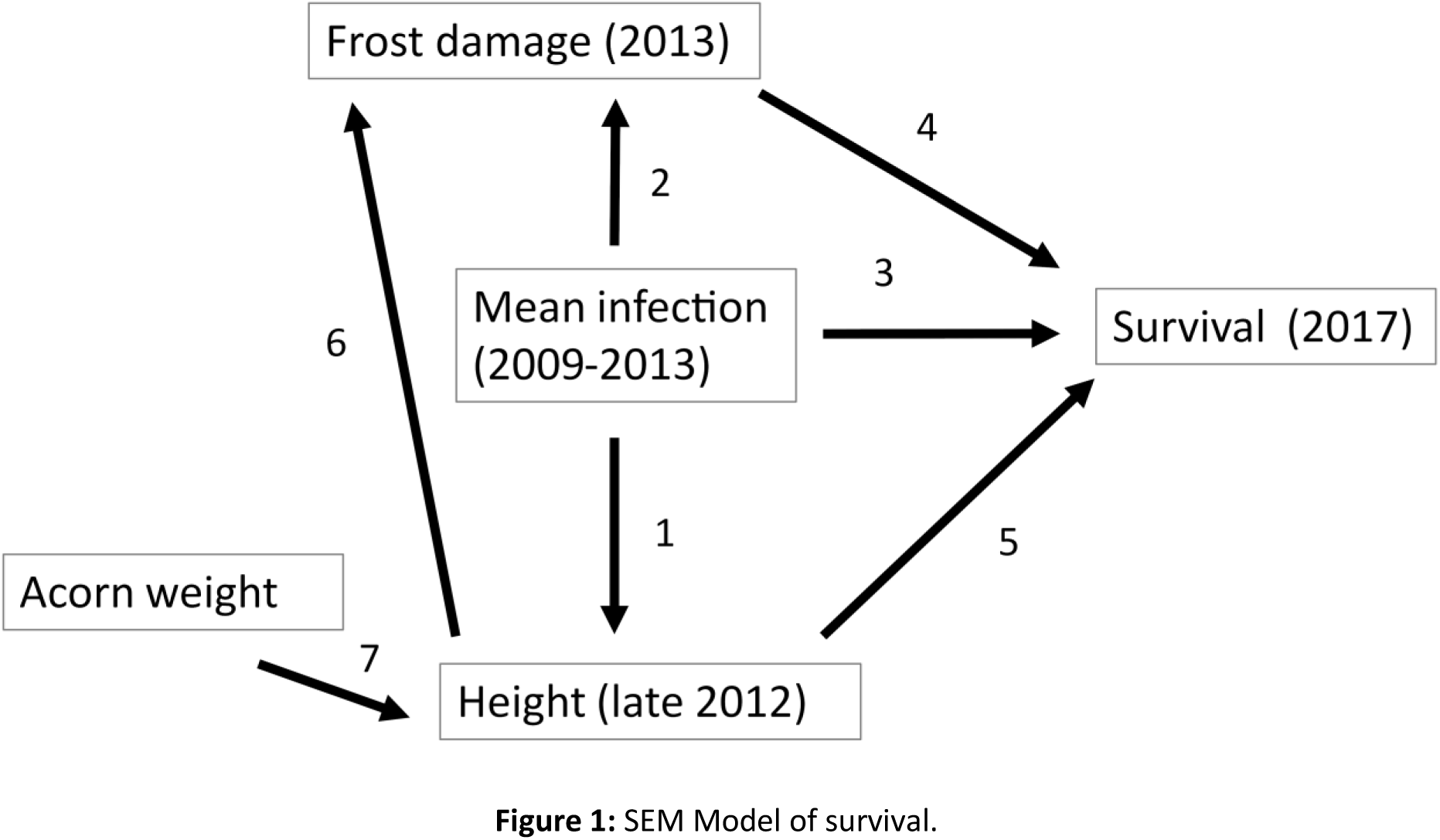
SEM Model of survival.

One path was an indirect effect of powdery mildew infection on survival through an effect on height. This was based on the assumption that infection has a direct (negative) effect on growth (as reported in Bert et al 2016), thus on height (arrow “1”) and that seedling height is expected to have a direct effect on survival (arrow “5”). The second path was an indirect effect of powdery mildew on survival through a direct effect on frost damage (“Frost damage 2013”, arrows “2”), and a direct effect of frost damage on survival (“Survival (2017)”, arrow “4”). This path is consistent with previous observations on the same field experiment (Desprez-Loustau et al 2014) or made by other authors (reviewed in Marçais & Desprez-Loustau 2014) that suggested that powdery mildew infection could affect the cold hardening process of shoots at the end of the season, resulting in greater shoot mortality during winter. A third path was a direct effect of powdery mildew infection on survival (arrow “3”) which may include toxic effects of the pathogen on its host or other effects not taken into account by the other paths. Finally, we included two other effects not related to powdery mildew infection: a direct effect of height on frost damage and one of acorn weight on height (arrows “6” and “7”, respectively).

#### Analyses at family level

Since the same 15 families were tested under both powdery mildew exposures (Natural *versus* Protected due to fungicide use), their relative performance in both environments could be compared based on family mean phenotypic value. In particular, the proportion of individuals having survived in each family is an estimate of one component of the reproductive success of their mother tree under each environment. Moreover, the family mean of each trait could be considered as an estimate of the genetic value of the corresponding mother tree. The relationship between family growth potential (*i.e.*, defined as the mean progeny height in a reduced disease environment provided by the Protected exposure) and family disease resistance (inversely related to mean progeny infection scores under natural conditions) can then be analyzed.

In order to assess temporal changes in the relative family composition of the surviving populations under both disease exposures, we calculated a Shannon Index in each plot and year as H’ = −∑p_i_*log_2_(p_i_), with p_i_ the proportion of each family in the population. H’ can vary between a maximum value of log_2_(N), with N the number of groups, if all groups have the same frequency (here N=15 and H’max=3.91), and a minimum value of 0 if the population is composed of a single entity.

All analyses were performed with the SAS software Version 9.4 (Copyright © 2013, SAS Institute Inc., Cary, NC, USA), in particular the Logistic, GLM and Calis (for SEM) procedures (scripts and data can be found in the zenodo repository https://doi.org/10.5281/zenodo.7517641).

### Genetic analyses

#### Sample collection and DNA extraction

Three to six leaves were sampled on each emerged seedling at the 15-leaf stage so as not to compromise the survival of individuals. Nine 9-mm-diameter leaf discs were cut off from the dried-leaves for each individual and stored at −80°C in 96-well plates until DNA extraction. DNA was extracted using the Invisorb DNA plant HTS 96 kit (Invitek, Germany). We followed the manufacturer instructions except that samples were disrupted with two 4-mm tungsten carbide beads during 2 X 1 min, at 30 Hz and that the lysis step lasted 1 hour (instead of 30 min) at 65°C. A Mixer Mill MM300 (Retsch, Germany) was used to disrupt the leaf samples. DNA was eluted in a final volume of 60 µl of elution buffer. SNP genotyping required high DNA quality and quantity. Genomic DNA samples quantity was assessed using the Quant-iT™ PicoGreen® dsDNA Assay Kit (Invitrogen™) according to the manufacturer’s instructions. Absence of DNA degradation was controlled on 1% agarose gel by the DNA bank platform of the Genotyping National Center, CNG (CEA-IG, Evry, France). A second genomic DNA extraction was performed for samples where concentration was lower than 45 ng.µl^-1^, or if the total amount of DNA was lower than 1 µg. Assignment of individuals to their half-sib families was checked using nine microsatellites (Guichoux et al 2011).

#### SNP selection and array design

The SNPs were chosen among a subset of 8,078 polymorphic SNPs from the allelic resequencing of more than 800 initial targeted genic regions within the genome of 13 *Q. robur*, using a high quality SNP database from Sanger sequence data, (Lang et al 2021; https://github.com/garniergere/Reference.Db.SNPs.Quercus). Two Perl scripts, *SNP_statistic* from the SeqQual pipeline (https://github.com/garniergere/SeqQual) and *snp2_illumina* (Lepoittevin et al 2010), were used to compute statistics for each SNP and to design a template file compatible with the Illumina Assay Design Tool (ADT) software respectively. Within the 13 *Q. robur* sequence data, three criteria were used to further filter SNP genotypes: a minimum depth of 8 reads, a minor allele frequency higher than 7% (allowing exclusion of variants found only once for the minimum of 8 diploid individuals sequenced) and an Illumina ADT score greater than 0.4, which yielded 2,447 SNPs. Moreover, in the case of two SNPs within 60 bp of each other, only one was kept, following Illumina’s recommendations, the chosen SNPs fulfilling the same previous quality criteria (Supplementary material, Figure S3), which yielded 1,670 SNPs. Finally, two stringent filters were added: (i) SNPs with an ADT score lower than 0.6 and with only 2 sequences for one of the alleles and (ii) SNPs with a minor allele frequency lower than 10%, with no heterozygous individuals identified and with only 2 sequences for one of the alleles. These two categories of SNPs were excluded. Finally, 1536 SNPs were included in the genotyping assay.

#### SNP Genotyping

The SNP genotyping experiment was performed on the subset of seedlings with the highest quality and quantity of extracted DNA, 1,185 individuals being finally retained (*i.e.*, 71% of those that underwent DNA extraction) with 759 and 426 individuals for the Natural and Protected exposure, respectively. For each 96-well plate, we checked the quality and reproducibility of the genotyping essay with one negative control (water) and four positive controls (DNA samples of two well-known genotypes, 3P and A4, duplicated twice). We also included across plates 59 DNA samples of parents and potential parents to further test for possible Mendelian inconsistencies between parents and offsprings. A total of 30-50 ng of genomic DNA per individual was used for SNP genotyping by the INRA-EPGV group using the Illumina BeadArray platform of the Genotyping National Center, CNG (CEA-IG, Evry, France) and following the GoldenGate Assay manufacturer’s protocol (Illumina Inc., San Diego, CA, USA). Three assays, over a 3-days period each, were performed to genotype 1,284 samples for the 1,536 SNPs. The protocol was similar to the one described by Hyten et al (2008), except for the number of oligonucleotides involved in a single DNA reaction, which comprised 4,608 custom oligonucleotides in the Oligo Pool Assay (OPA). Raw hybridization intensity data processing, clustering and genotype calling were performed using the genotyping module of the BeadStudio/GenomeStudio package (Illumina, San Diego, CA, USA) with a GeneCall score cutoff of 0.25 to obtain valid genotypes for each individual at each SNP.

#### SNP quality criteria for genotyping reliability

After a first genotype calling of the raw data, we assessed SNP genotype quality across individuals using the methodology proposed by Illumina (Tindall et al 2010). Briefly, 50% GC score and 10% GC score were plotted as a function of the sample call rate. Poorly performing samples were obvious outliers across many genotypes when compared to the majority. In our experiment, these outliers corresponded to samples with 50% GC score and call rate lower than (mean(50% GeneCall score)-0.0.1) and (mean(call rate)-0.015) respectively or to samples with 10% GeneCall score and call rate lower than (mean(10% GeneCall score)-0.0.15) and (mean(call rate)-0.015) respectively (Figure S4). After discarding those poor-quality samples, a new genotype calling was performed on remaining individuals using the same GeneCall score cutoff. SNP quality was further determined automatically using a call frequency greater than 0.99, a 10% GeneCall score greater than 0.6, a heterozygote frequency greater than 1%, and a very low level of inconsistencies for Parent-Child or Parent-Parent-Child testing. To avoid discarding valuable SNPs or keeping poor quality SNPs, a visual inspection of all SNPs clusters was further performed after the automatic pipeline. SNP markers that displayed either compression or unexpected clustering patterns were discarded (Supplementary material, Figure S5 and S6). A total of 819 SNPs were finally kept for further analyses (Supplementary material, Table S3).

#### Multi-locus individual genetic diversity

Genetic diversity indices were computed for different groups of individuals: the 1185 individuals that were representative of the initial populations, and individuals that survived or not under both powdery mildew exposures (Natural and Protected). Observed and expected heterozygosities (H_o_ and H_e_, *sensu* Nei 1973), and F_ST_ indicating differentiation between the initial and the surviving populations for both exposures were estimated (Weir and Cockerham 1984), using ‘adegenet’ (Jombart 2008) and ‘Genepop’ (Rousset et al 2008) R packages. Differentiation between populations and deviation of populations from Hardy-Weinberg equilibrium were tested using ‘Genepop’ R package (Rousset et al 2008). Five heterozygosity statistics were estimated for each individual based on the 819 successfully genotyped SNPs, using the GENHET function in R (Coulon 2010): the proportion of heterozygous loci (PHt), two standardized heterozygosities based on the mean expected heterozygosity and on the mean observed heterozygosity (Hs_exp and Hs_obs, respectively), the internal relatedness (IR) and the homozygosity by locus (HL). The preliminary analyses showed that these five statistics were highly correlated (absolute Spearman’s rank correlation coefficient between 0.96 and 1.00; Supplementary material, Figure S7). Therefore, only PHt was kept for further analyses. Mean estimates of PHt were compared among the group of individuals that did survive *versus* the ones that did not for each exposure (Natural and Protected) using Mann-Whitney tests. Equality of variances were also assessed using Fligner-Killeen tests among the same group of individuals (survivors *versus* dead individuals, both for Natural and Protected exposures). The mean values of PHt in Natural *versus* Protected, initial and surviving, sub-populations were compared by running a GLM model. The effect of PHt on survival was also tested by logistic regression along with four other main explanatory variables (Supplementary material, Table S2, Model 6).

#### GWAS analysis

The physical position of SNP markers was obtained by aligning the flanking sequence of the SNP markers (with a maximum of 100 base pairs on each side of the SNP; Data S1) using Blast (Johnson et al 2008) on the *Q. robur* genome assembly available on The Darwin Tree of Life project (accession PRJEB51283). This confirmed that the 819 SNPs were scattered across the whole genome, on all chromosomes, in 426 different genic regions (see Table S3), with an average distance between regions of 2.11 Mb (ranging from 0.0029 to 15.3 Mb). The associations between SNP markers and the four phenotypic traits of interest (“Mean infection (2009-2013)”, “Height in 2012”, “Acorn weight” and “Survival (2017)”) were tested using the Bayesian-information and Linkage-disequilibrium Iteratively Nested Keyway (BLINK, Huang et al 2019) in GAPIT3 R package (Wang and Zhang 2021). Default parameters were used for all analyses. This method allows to account for the different relatedness levels among individuals, building from the multilocus mixed model of Segura et al. (2012) by iteratively incorporating associated markers as covariates, but with a special optimization criterion (Tibbs Cortes et al 2021). Because of the large number of tests, a false discovery rate (FDR) analysis was used to control for false positive associations (Benjamini & Hochberg 1995), using a threshold of 0.01 for the FDR-corrected p-value. Deviation of the observed p-values from the expected values was assessed with a QQ-plot (Supplementary material, Figure S8 and S9 for Natural and Protected exposures, respectively). Both powdery mildew exposures were analyzed separately.

## Results

### Seedling and juvenile survival

Oak survival was very high during the first five years, close to or greater than 90% in both powdery mildew exposures, *i.e.*, the protected or fungicide-treated one, and the natural or non-protected one that was submitted to natural powdery mildew infection (Figure 2A). Survival decreased in the following four years. The decrease was much steeper for non-protected trees. Mortality was observed mainly at the beginning of spring when individuals failed to flush, and not during the growing seasons. The annual mortality rate was highest in 2014 in all non-protected plots but one, compared to the fungicide-treated plots. Survival at the end of the monitoring period was 49.7% on average in plots with natural powdery mildew infection, compared to 79.5% in fungicide-protected plots. As expected for an efficient fungicide treatment, disease severity was much lower in fungicide-treated plots than in non-protected plots throughout the experiment (Figure 2B). Seedlings showed higher height in the protected plots than in the naturally infected plots, as soon as the second year (Figure 2C).

**Figure 2:**
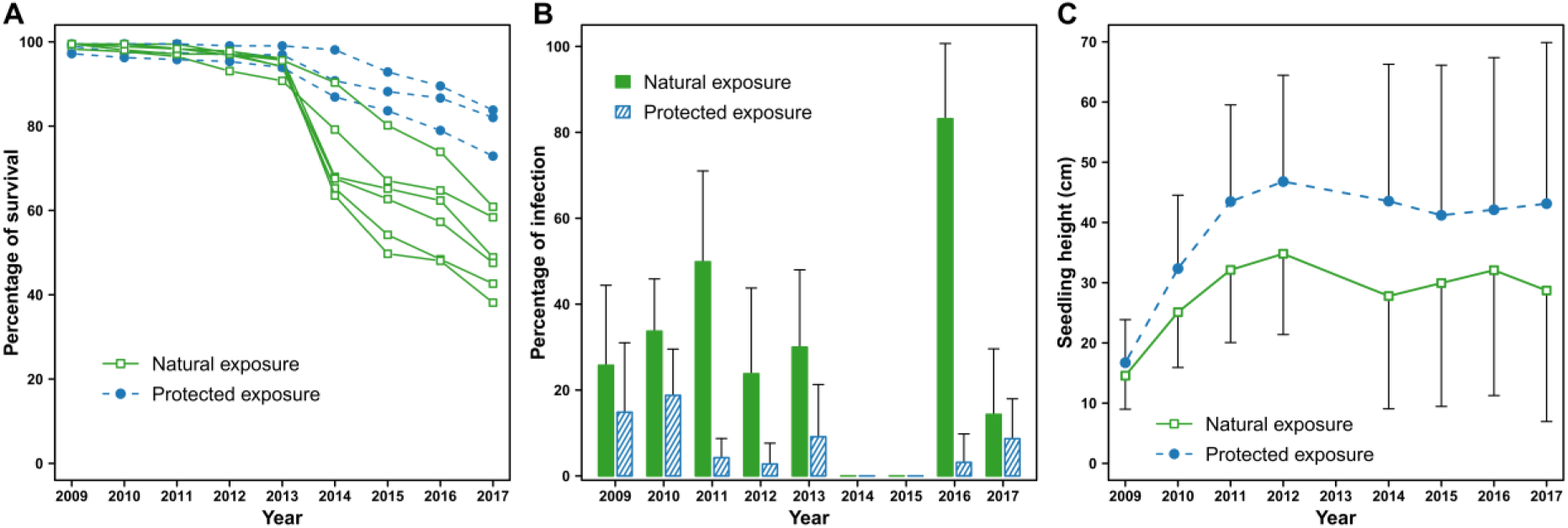
A. Time course of individuals’ survival rate in the three protected replicates and the six natural infection plots; B. Mean annual infection across plots under both powdery mildew exposures; C. Seedling height across studied years in protected *versus* natural exposure plots.

The powdery mildew exposure and the acorn weight were both significant predictors of survival in the last year of observation *i.e.*, 2017 (Wald Khi2 = 4.00, P = 0.045 and Wald Khi2 = 27.1, P < 0.0001, respectively; Supplementary material, Table S4 and Figure S10). The natural mildew exposure was associated with a four-fold increase in the odds of mortality compared to the fungicide-protected one, which corresponds to an odds ratio of 4.0 with a 95% confidence interval (C.I.) of 3.19 to 5.06. For the acorn weight, a 23% increase of the odds of survival per additional gram was observed on average (odds ratio of 1.23 with a 95% C.I. of 1.14 to 1.32). The interaction effect between acorn weight and powdery mildew exposure was not significant in the model (Supplementary material, Table S4), as well as the block effect (not shown in the final analysis).

Model 2 provided a quantitative assessment of the effect of the mean percentage of infection across the first five years, on survival (Supplementary material, Tables S2 and S5), with an estimated odds ratio of 0.954 (95% C.I of 0.946 to 0.962). This means that each additional percent of leaf infection is expected to reduce the odds of survival by 4.6% (Figure 3).

**Figure 3:**
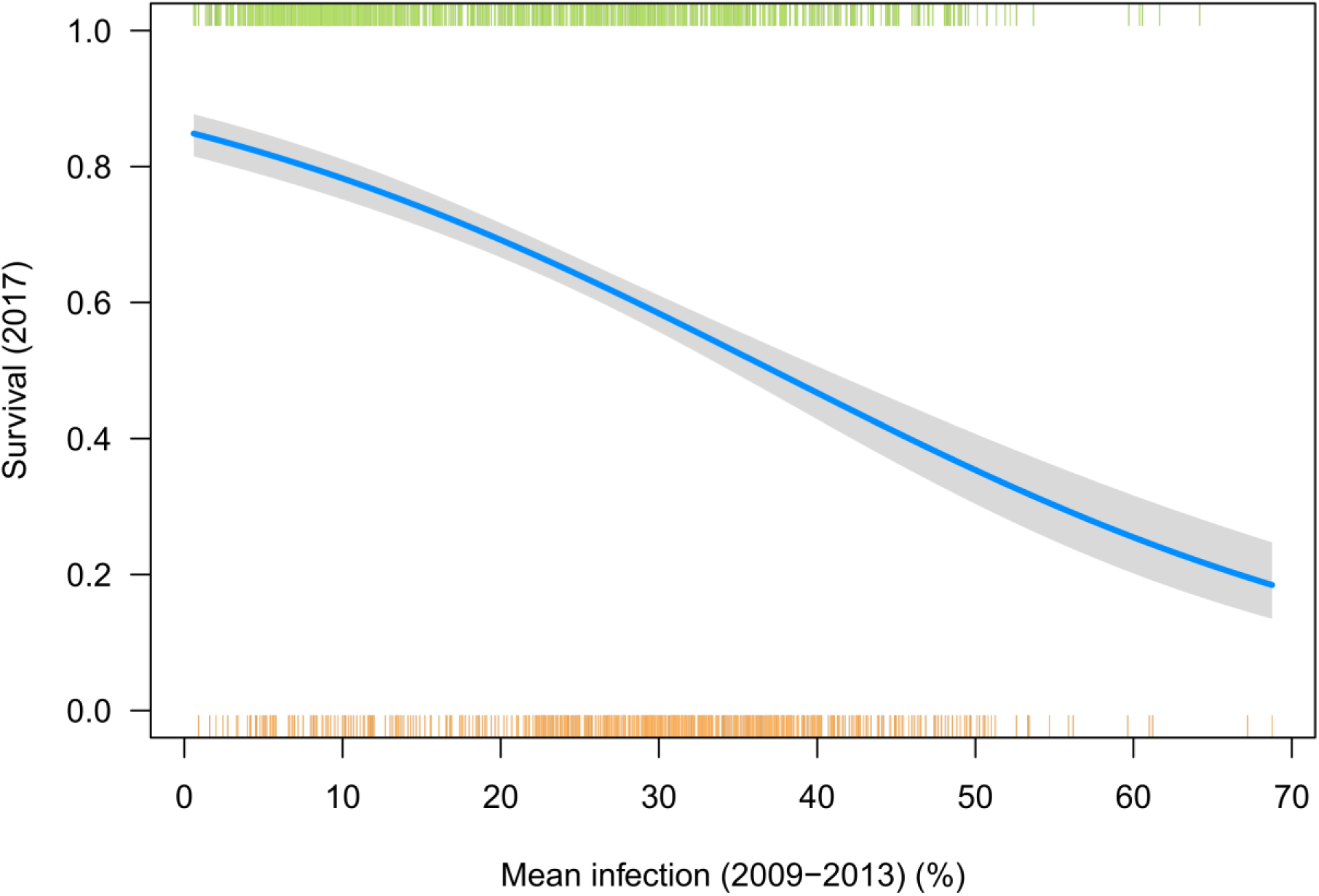
Logistic model predicting juvenile survival in year 2017, based on Mean infection between 2009 and 2013 (at mean acorn weight = 5.028 g). Distributions of the individual values of the variable “Mean infection (2009-2013)” were displayed as orange marks at the bottom of the figure and light-green marks at the top of the figure for dead and live trees, respectively. The grey envelop around the blue line represents the 95% confidence interval.

Using seedling height at the end of the first growing season (Model 3, Table S2) instead of acorn weight as a predictor variable (Model 1, Table S2) had little influence on the results, with a very slight improvement of the concordance of the association between predicted probabilities and observed responses (68.7% instead of 68.1%). Thus, seedling height at the end of the first growing season was a good predictor of survival at the end of the monitoring period (*i.e.*, eight years later), with a strong negative impact of the “Powdery mildew exposure” at a given height (Supplementary material, Figure S11). The “Height in 2009 : Powdery mildew exposure” interaction was not significant in this model. When “Frost damage (2013)” was added to the logistic model (Model 4), this variable had a significant negative effect on survival (Wald Khi2 = 5.99, P = 0.014, odds ratio = 0.804; C.I. of 0.649 to 0.996) in addition to the effects of “Acorn weight” and the “Mean infection (2009 and 2013)” (Wald Khi2 = 17.8, P < 0.0001 and Wald Khi2 = 112.8, P < 0.0001, respectively; Supplementary material, Figure S12).

The Structural Equation Model showed almost equal but opposite effects of “Height (late 2012)” and “Mean infection (2009-2013)” onto final survival, with total standardized coefficients (not displayed on Figure 4) of 0.30 (positive) and −0,28 (negative), respectively. The total negative effect of powdery mildew infection corresponds to a direct negative effect of −0.19 (path 3) and indirect negative effects of −0.09. The most important indirect effect is −0.087 (=-0.28*0.31, according to paths 1 and 5) through “Height (late 2012)”, the indirect effect through “Frost damage (2013)” being much less (0.08*-0.09=-0.007, according to paths 2 and 4) (Figure 4). The direct contribution of “Frost damage (2013)” on final survival (path 4) was a mild negative effect (−0.09) (Figure 4).

**Figure 4:**
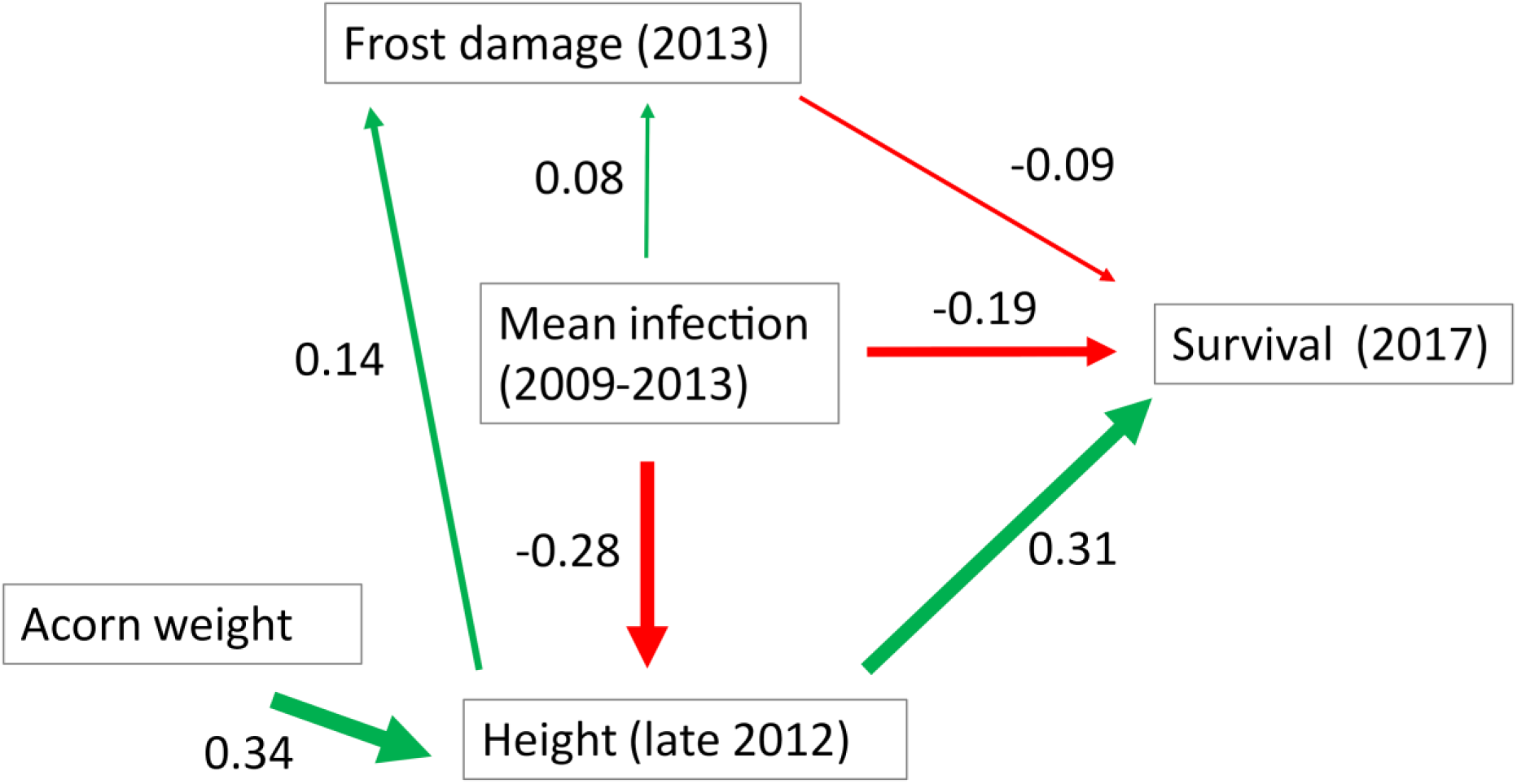
Results of the Structural Equation Model – Line width is proportional to the effect value.

### Differential impact of powdery mildew among open-pollinated families?

Average proportions of individuals having survived varied among families, ranging from 35% to 93% in the fungicide-protected plots and from 23% to 72% in the plots submitted to natural powdery mildew exposure (Figure 5).

**Figure 5:**
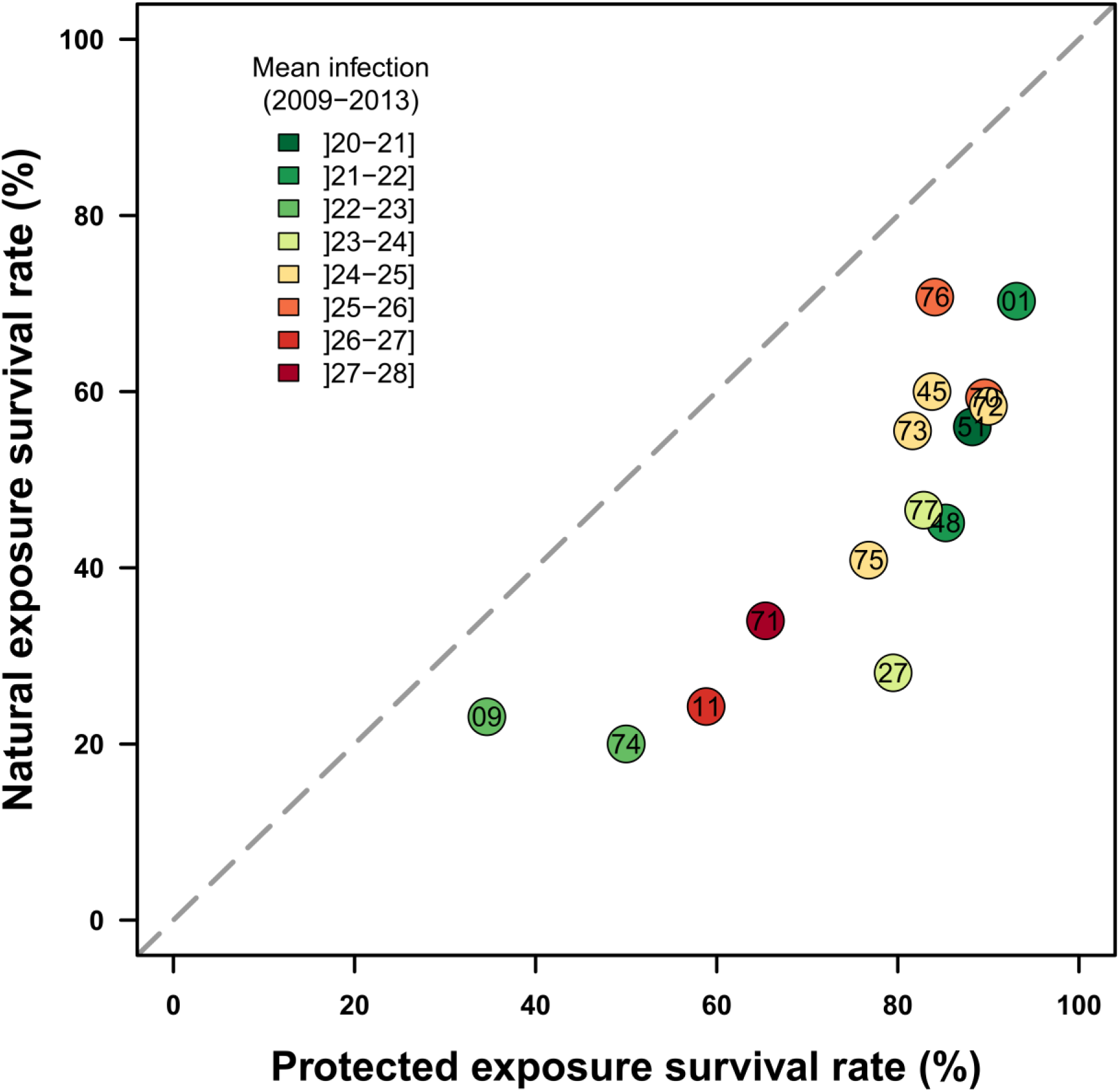
Progeny survival percentages across families in Protected plots *versus* Natural powdery mildew infected plots. Each dot corresponds to a family identified by its number. The mean powdery mildew infection (2009-2013) for each family in the Natural infection exposure is color-coded from dark-green = low mean powdery mildew infection (minimum=20.9) to dark-red = high mean powdery mildew infection (maximum=27.8).

The full logistic model of survival, including previously studied factors (powdery mildew exposure, acorn weight and frost damage), a family effect and an interaction between powdery mildew exposure and family (Model 5) was highly significant and showed an improved concordance percent between predictions and observations of 76.7%. All previously studied factors remained significant but the family effect further explained the probability of survival (Wald Khi2 = 126.68, P < 0.0001; Supplementary material, Table S6).

However, the interaction between family and powdery mildew exposure was not significant. This means that overall, in this experiment, exposure to powdery mildew had a similar negative effect on the survival of all families without any strong changes in their ranking for survival (Spearman correlation = 0.86; P = 0.0001, and see Figure 5).

Mean family survival (percent surviving progeny in 2017) was significantly correlated with family height (*i.e.*, mean value over the progeny) from 2014 onwards in both disease exposures (*e.g.*, r = 0.71 and 0.69 with height in 2017 in fungicide-protected and powdery mildew natural exposures, respectively). In plots under the natural exposure, the relationship between family survival and family “height potential”, *i.e.*, mean height of the same family measured in protected plots in 2017, was even stronger than with realized height (r = 0.82, P = 0.0002). No significant correlation was observed at family level between height potential (in any year) and powdery mildew susceptibility (= mean infection observed in powdery mildew exposed plots in 2009-2013), although both variables showed a significant family effect. The height of surviving juveniles at the end of the monitoring period (in 2017) varied significantly among families: from 19.2 cm to 53.4 cm in fungicide-treated plots (F = 5.01 – df = 14 P < 0.0001), and from 17.8 to 35.7 in powdery mildew plots (F = 2.38 – df = 14 P < 0.0032) (Figure 6).

**Figure 6:**
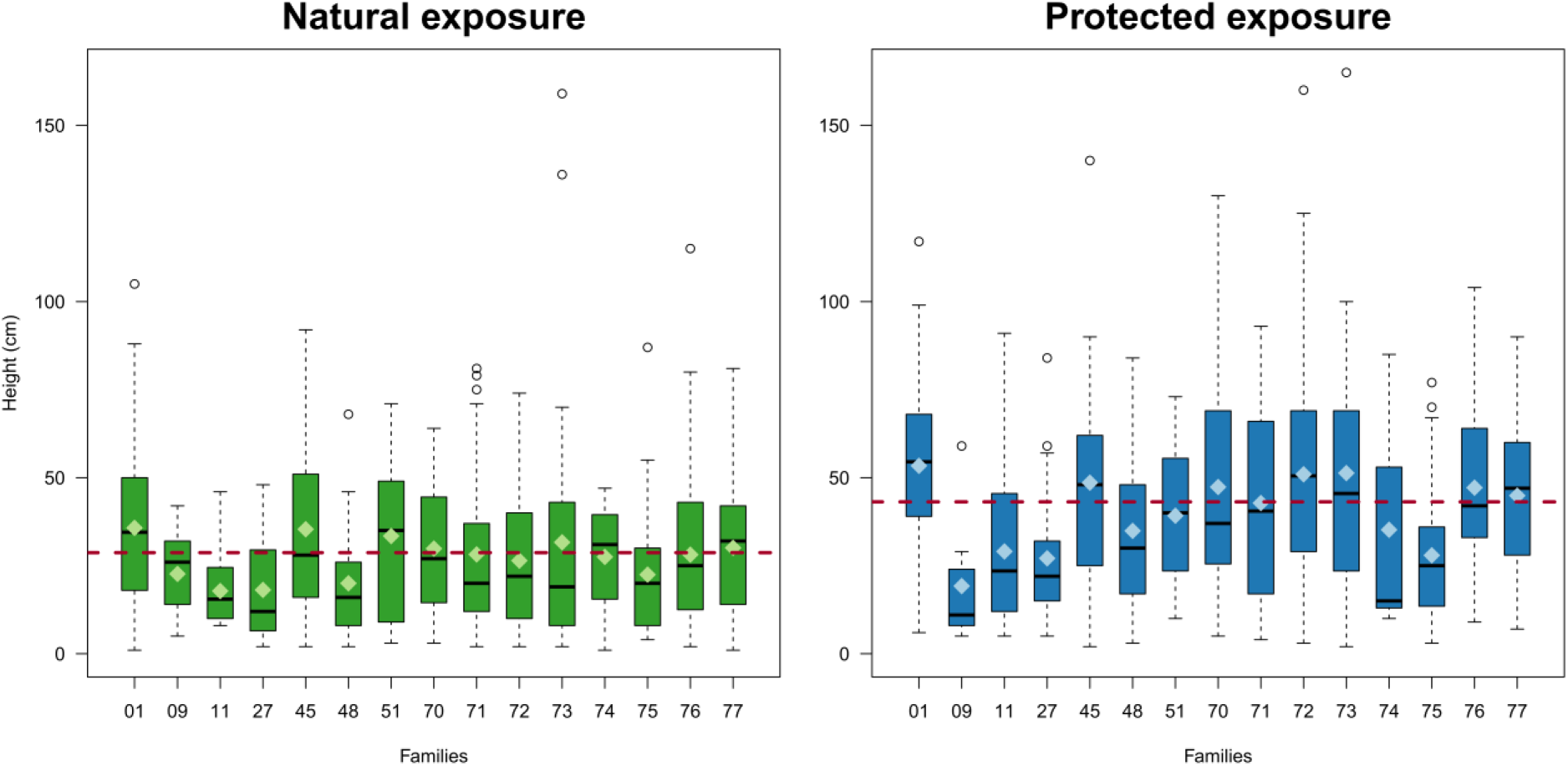
Boxplot of recorded heights in 2017 in both Natural (left) and Protected (right) exposures across families. The black lines represent the median height for each family and the light-colored diamonds represent their mean height. The red dotted lines represent the overall averages of juvenile height in each exposure.

The between-family coefficient of variation for final height (standard deviation/mean) was lower in the natural disease exposure than in the protected by fungicide exposure (21.5 and 26.4, with standard deviations = 5.84 and 10.54, respectively) (Figure 6). Within-family variation (SD) was also reduced in the natural disease exposure compared to the protected exposure (t = −2.4; P = 0.0306). Mean powdery mildew infection over the first five years under the naturally exposed plots varied significantly among families from 28.5 to 35.7% (F = 5.28 – df = 14 P < 0.0001).

The Shannon index, measuring diversity within plots in terms of family composition, remained very high in fungicide-treated plots throughout the experiment, but decreased after 2013 in all plots naturally exposed to powdery mildew, as a result of increasing differences across years in the relative numbers (percent of surviving individuals) of the different families (Figure 7).

**Figure 7:**
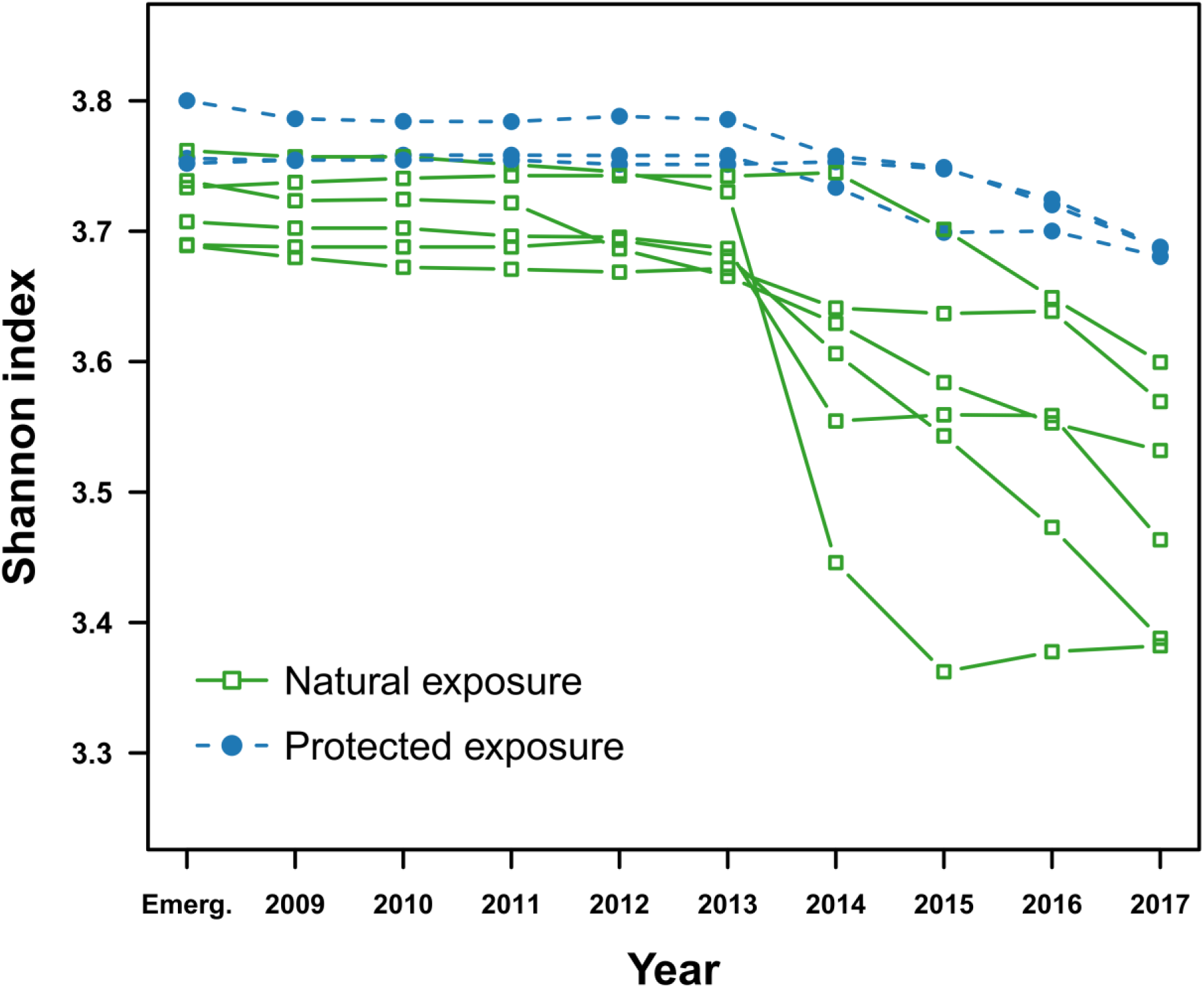
Temporal changes in Shannon index, used as a family-diversity index calculated from the proportional family abundances in plots under natural or protected by fungicide powdery mildew exposure. The index was calculated for each of the nine experimental plots: six under natural powdery mildew exposure (green) and three under protected by a fungicide powdery mildew exposure (blue) every year.

### Multi-locus heterozygosity

Out of the 1,185 individuals included in the SNP genotyping experiment, 1,143 were successfully genotyped with less than 0.08% of missing data. Observed and Expected heterozygosity did not vary between initial and surviving populations in both disease exposures, with values of Ho and He of 0.32-0.33 in all cases (Supplementary material, Figure S13). Genetic differentiation between initial and surviving populations were very low and not significant (Supplementary material, Figure S13). The distribution of the proportion of heterozygous loci (PHt, see methods) values estimated on all individuals was negatively skewed: while most individuals had a PHt value in the range of 0.235-0.335, very few individuals (2.54%) showed lower values (< 0.235, minimum 0.18) (Supplementary material, Figure S14). The mean PHt across individuals was very similar in both disease exposures when comparing initial *versus* surviving populations: 0.283 ± 0.022 *versus* 0.284 ± 0.020 for the natural exposure; 0.283 ± 0.025 *versus* 0.285 ± 0.019 for the protected exposure (Supplementary material, Figure S15 and Table S7). However, in both exposures and across all families, the individuals with very low PHt were over-represented in dead seedlings (Figure 8, Supplementary material, Figure S16). This resulted in a decrease in the variance of the PHt between surviving and dead individuals (Khi2 = 0.567, P = 0.45 and Khi2 = 11.4, P = 0.0007 for Natural and Protected exposure, respectively).

**Figure 8:**
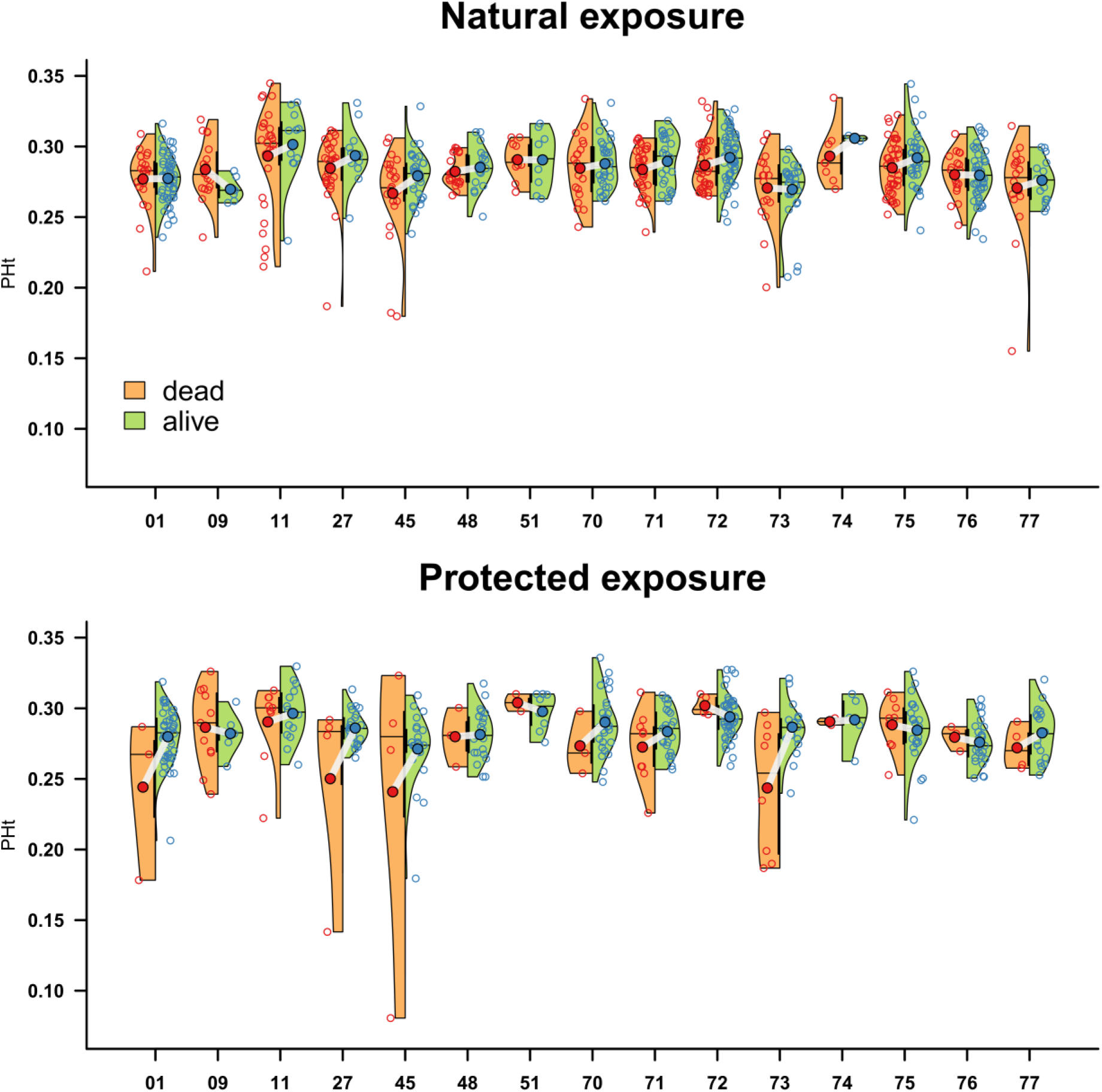
Violin plots of the PHt values for Natural (above) and Protected (below) exposures across families. Red and blue empty (or full) circles represent the PHt values, across dead and alive individuals (or families), respectively. Black horizontal lines delimit the median values for dead and alive individuals across families.

The logistic model of survival including PHt in addition to the exposure (Natural *versus* Protected), family, acorn weight and frost effects (Model 6, Table S2) demonstrated a significant positive effect of individual heterozygosity on survival, but the Powdery mildew exposure*PHt interaction was not significant. This suggests that the effect of low heterozygosity was not more deleterious in naturally exposed than in protected seedlings but simply added to the negative effect of infection.

### Tests of genetic associations

Overall, 16 significant genetic associations were found between 14 loci (SNPs) and the four phenotypic traits investigated, mostly on chromosomes 2, 6 and 8 (Figure 9). In the Natural exposure, one SNP was statistically associated with “Mean infection (2009-2013)” on chromosome 6. This SNP belonged to the same gene as another close one (distant from 336 bp) that was significantly associated to “Acorn weight” (Figure 9). One SNP located on chromosome 8 was associated with “Survival (2017)”. Two SNPs were significantly associated with “Height in 2012”, one of which was also associated with “Acorn weight” (Figure 9). In the protected exposure, a different SNP located on chromosome 2 was associated with “Survival (2017)”. This SNP was also associated with “Acorn weight”.

**Figure 9:**
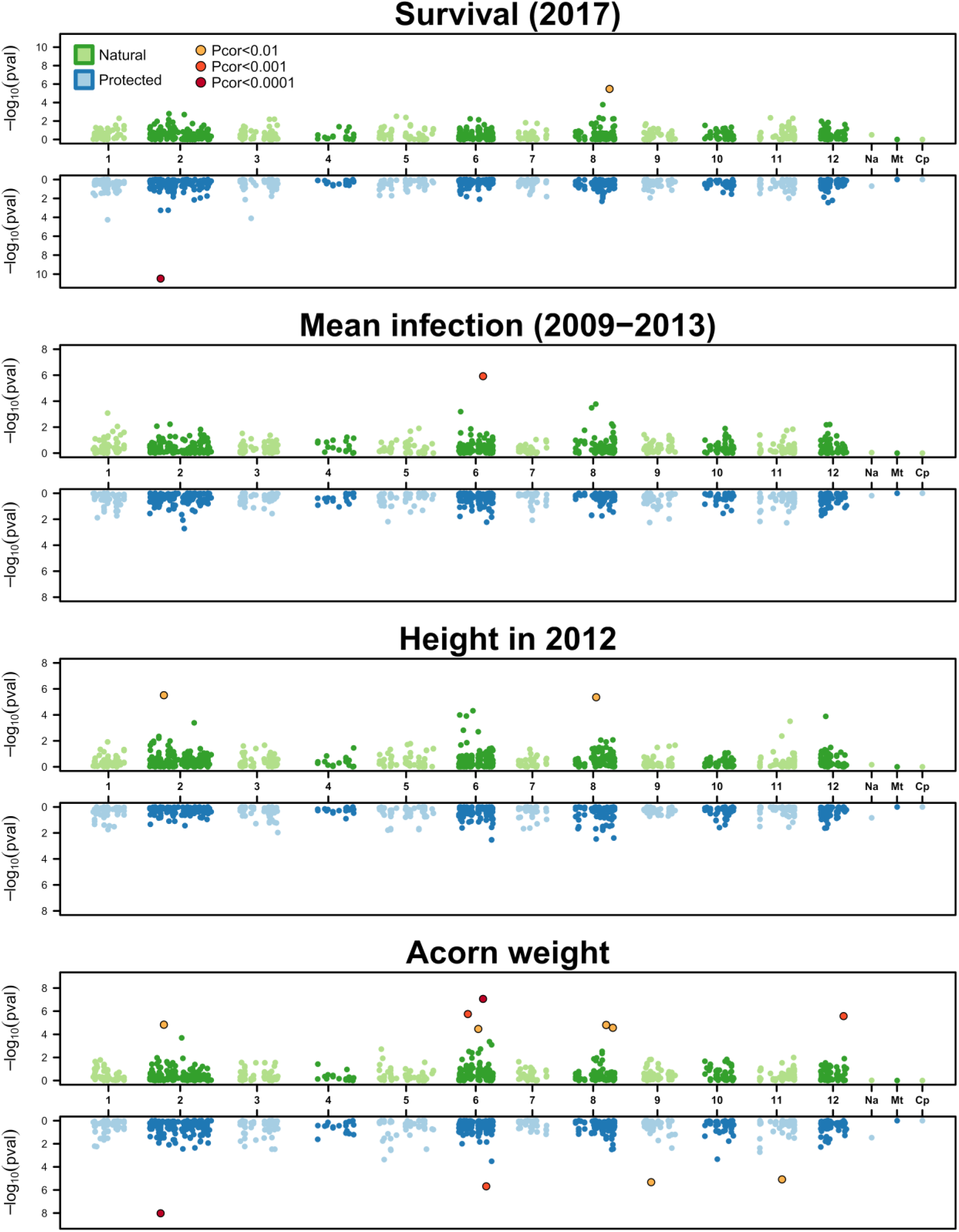
Manhattan plot for the Genome Wide Association Study results across the four phenotypic traits investigated: “Survival (2017)”, “Mean infection (2009-2013)”, “Height in 2012” and “Acorn weight”, and across both exposures (Natural in blue, Protected in green). Each dot represents a SNP. The negative logarithm of the association *p*-value corrected for multiple tests is displayed on the vertical axis. SNPs with a significant association with a trait are indicated with yellow, orange and red dots. The SNP markers are ordered along the genome and grouped by chromosome. *1* to *12*: chromosome number; *Na*: unknown location; *Mt*: mitochondrial; *Cp*: chloroplast.

The SNP *CL7647CT8535_01-89* linked to “Mean infection (2009-2013)” is located in a gene predicted to be an ethylene response factor C3. While no significant association of this SNP was found with seedling survival in the natural exposure, the genotypic classes with lower mortality are consistent with those showing less infection and thus increased resistance (Figure 10, top row). “Acorn weight” and “Height in 2012” did not show any association with this marker. The SNP *CL8450CT11856_03_04-703* linked to survival in the natural exposure is located in a gene coding for a putative histone H4. No differences were observed for the three other traits among genotypic classes at this locus (Figure 10, middle row). The *SNP CL8754CT10139_03-29* significantly associated with the survival in the protected exposure is situated in a gene coding for a putative pentatricopeptide repeat-containing protein. This marker was also significantly associated with “Acorn weight”, but not with “Height in 2012” or “Mean infection (2009-2013)” traits (Figure 10, bottom row).

**Figure 10:**
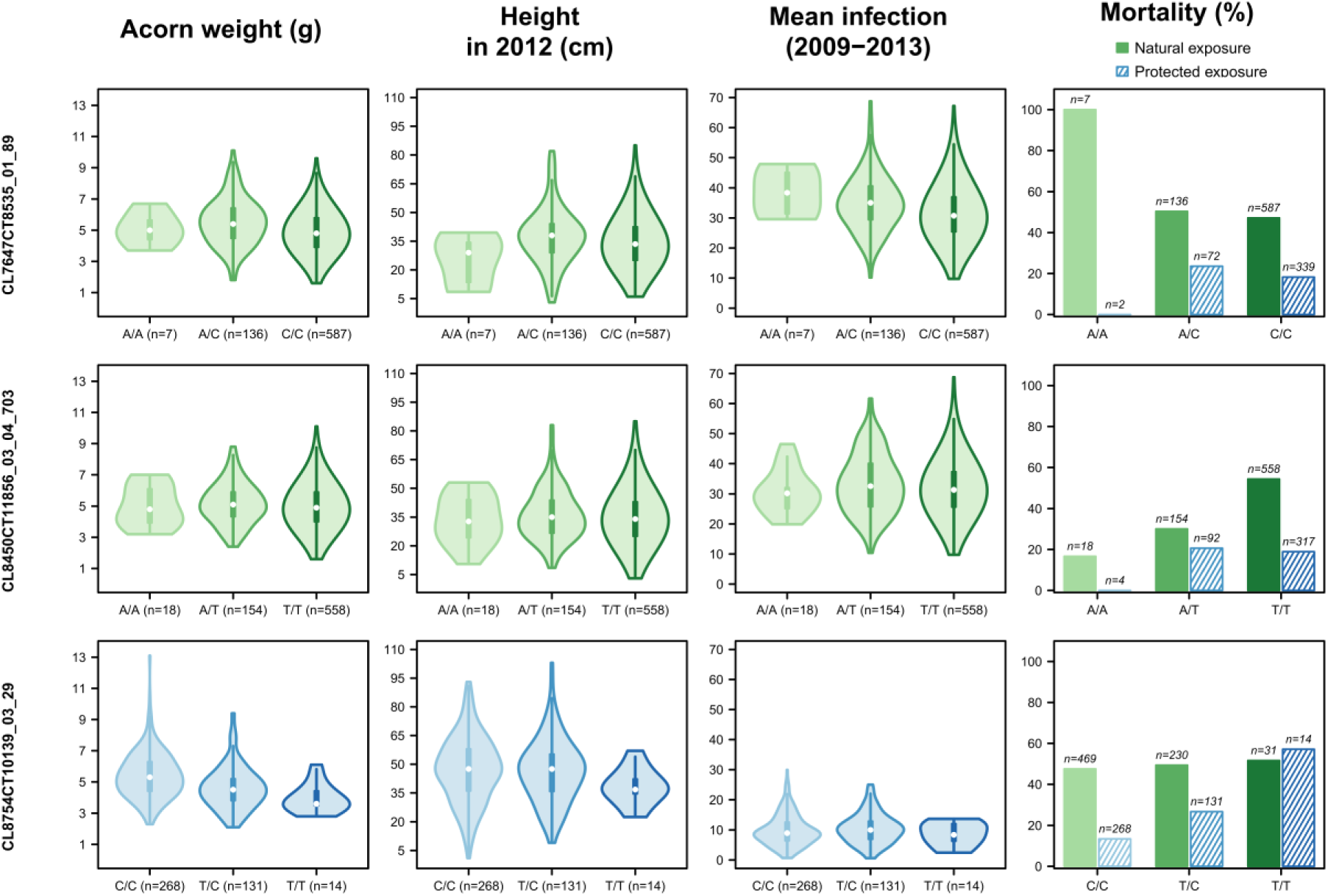
Genotypic classes distribution and mean trait values across traits for SNPs *CL7647CT8535_01-89*, *CL8450CT11856_03_04-703* and *CL8754CT10139_03-29* that were significantly associated with “Mean infection (2009-2013)” under natural exposure, “Survival (2017)” under natural exposure and “Survival (2017)” under protected exposure (last column), respectively. Genotypic classes are named using the IUPAC nucleotide symbol convention: A= adenine, C=cytosine and T=thymine.

## Discussion

In this experimental approach, the effect of powdery mildew on oaks was monitored during nine years after acorn sowing in contrasting plots that were either under natural infection or where pathogen pressure was limited by fungicide treatments. More than one thousand individuals identified by their acorn weight and mother tree (15) were subjected to fine phenotypic monitoring across years (height growth, mildew infection and survival) and genotyped at 819 SNPs. This powerful design enabled us to demonstrate various effects of powdery mildew at demographic and genetic levels in early stages of an oak cohort. The significance of these results and a number of related issues are discussed below.

### 1. Strong negative powdery mildew impact on juvenile survival

The negative impact of powdery mildew on oak regeneration was pointed out by many authors (Pap et al 2012; Marçais & Desprez-Loustau 2014; Demeter et al 2021). However, we could not find any quantitative data on pathogen-induced mortality of seedlings in forest. Our experimental approach, with a comprehensive individual monitoring of seedlings across nine years under two contrasted disease exposures, provides supporting evidence of a causal association between powdery mildew infection and mortality of seedlings, under field conditions. Mortality was indeed significantly much higher in the plots exposed to natural infection than in plots treated with fungicide. Moreover, the probability of mortality could be quantitatively related to disease severity in the previous years.

The high mortality rates in early stages of naturally regenerated forest stands are generally attributed to an intense competition among tree seedlings (Collet & Le Moguedec 2007). However, mortality patterns in our experiment suggest that powdery mildew effects overcame competition effects. Indeed, mortality rates remained very low in the fungicide-treated plots during the monitoring period even though plants were taller and maintained at a greater density than in plots without fungicide (where seedlings progressively died), thus at a potentially stronger competition level. Maybe the competition-related mortality (self-thinning stage) (Peet and Christensen 1987; Collet & Le Moguedec 2007) will simply be delayed in our conditions, characterized by full light availability and an initial seedling density (1 acorn per 10*10 cm) which may be lower than in some spots of natural regeneration (Diaci et al 2008; Annighöfer et al 2015; Kuehne et al 2020).

Infection induced mortality has been reported for other powdery mildew diseases, such as *Podospahera plantaginis* on *Plantago lanceolata* (Laine 2004), or *Erysiphe cruciferarum* on *Alliaria petiolata* (Enright et al 2007), and rust diseases (other plant biotrophic pathogens), such as myrtle rust (Carnegie et al 2016), *Melampsora medusa* f. sp. *deltoidae* on Poplar (Newcombe et al 1994) or *Puccinia lagenophorae* on groundsel (Paul & Ayres 1986). Mortality started only five years after sowing in our experiment, which suggests cumulative and delayed effects of infection, as expected for this kind of pathogen. As biotrophic parasites, powdery mildews strongly affect the carbon economy of their host plant, by direct consumption of carbon fixed by photosynthesis (through their haustoria) but also by forcing allocation of plant carbon to defense (Hückelhoven 2005, Oliva et al 2014). In addition, powdery mildew infection has a direct negative effect on net carbon assimilation by photosynthesis, as was demonstrated for *E. alphitoides* (Hewitt & Ayres 1975; Hewitt & Ayres 1976; Hajji et al 2009; Pap et al 2014). The depletion of carbon by the pathogen likely explains growth reduction. Cumulative and delayed effects of powdery mildew were previously described on radial growth in young oak trees (Bert et al 2016). Then it is reasonable to assume, although the full demonstration remains to be made (Martinez-Vilalta 2014), that severe infections, recurring in successive years, can lead to exhaustion of reserves and ultimately death

(Oliva et al 2014). The structural equation model that we tested is consistent with a strong direct effect of powdery mildew infection on seedling survival, twice as important as the effect mediated by decreased height. One possible mechanism could be reduced root growth in infected plants resulting from the alteration of the carbon metabolism. In the case of oak, the development of a large root system facilitates survival when aerial parts are affected or killed (Larsen & Johnson 1998). Finally, the SEM also suggested an indirect effect of powdery mildew through frost sensitivity, in agreement with previous observations of severe shoot mortality following winter in infected seedlings (Desprez-Loustau et al 2014). The late spring frost of 2013 could have given the “coup de grâce” to already weakened seedlings. Paul & Ayres (1986) also reported that heavy infection could compromise the ability of plants to tolerate winter stress in groundsel infected by rust. Jarosz & Burdon (1992), with flax rust, noted that the main effect of disease was to reduce survivorship during the winter following infection which could lead to pathogen-generated cycles in the host population size (Susi et al 2017).

### 2. Differential impact of powdery mildew across families

The observations of progenies from identified mother trees allowed to assess fitness components that were either linked to growth, disease resistance, or progeny survival of the mother trees that were originally sampled. Powdery mildew infection was quite high across years in our experiment and families showed different levels of disease severity (% leaf area infected). The progenies of all 15 mother trees were negatively affected in their survival under higher disease pressure. However, the ranking of the mean family values for survival was very similar under both powdery mildew exposures.

In particular, progenies from most competitive mother trees (*i.e.,* with greatest progeny survival under low disease pressure, with fungicide) were also among those with greatest survival under high powdery mildew pressure. The hypothesis of changes in mother tree survival ranking related to powdery mildew exposure was therefore not supported. This hypothesis was based on the assumption of a negative relationship between resistance to powdery mildew and growth (considered as an important component of fitness at seedling stage). In our experiment, seedling survival was indeed strongly related to growth, as estimated by seedling height. However, the results suggested there was no apparent trade-off between growth and disease resistance: the families with the greatest mean height in the fungicide-treated plots (*i.e.*, representing the growth trait) did not have the highest infection rates when exposed to disease (representing the defense trait). In both disease environments, the families with highest survival rates were also those with the greatest height growth potential (assessed under fungicide treatment).

Some features of our experiment may explain such absence of negative correlation between growth and disease resistance. First, only 15 mother trees were sampled on a small spatial scale in one local population, thus limiting the phenotypic variation that could be observed. It has to be noted however that genomic diversity was high, in the range of values, or greater than He values reported with the same type of markers (*i.e.* SNPs) for various *Q. robur* stands in Europe (Blanc-Jolivet et al 2021, Degen et al 2021 a and b). Trade-offs between traits (including disease resistance) may be easier to detect when considering phenotypic variation across a wider spatial range, in relation to differing selection pressures and evolutionary strategies of populations (McKnown et al 2014; Heckman et al 2019). In addition, the expression of growth-defense trade-offs can be context dependent (Karasov et al 2017), and it is usually stronger in environments where the level of resource acquisition is limited by shade or abiotic stresses (van Noordwijk & de Jong 1986; de Jong 1995).

Finally, the detection of growth-defense trade-offs may depend on the choice of the traits that are assessed. In our study, with height as the growth variable, we considered trade-off in a very general sense, encompassing processes linked to both acquisition and allocation of resources (Laskowski et al 2021).

Our results could also suggest that tolerance was more important than resistance to explain mean survival differences of progenies across mother oak trees under high disease pressure. Plants use different lines of defense to respond to pathogens, including resistance *sensu stricto* and tolerance (Desprez-Loustau et al 2016; Pagan & Garcia-Arenal 2020). Resistance *sensu stricto* relates to mechanisms that limit pathogen development within the plant. The variable corresponding to percent leaf area infected in our monitoring can be considered as inversely related to resistance. By contrast, tolerance relates to mechanisms with no direct effect on the pathogen but that limit the negative impact of infection on plant fitness (Jeger et al 2006).

We previously demonstrated that mechanisms such as increased polycyclism and compensatory growth are likely involved in the response of oak seedlings to powdery mildew (Desprez-Loustau et al 2014). In our experiment, mean survival across families in plots naturally exposed to powdery mildew, *i.e.*, one component of tree fitness, was not correlated with mean leaf area infection but significantly correlated with mean progeny height in protected plots (height potential). We can thus hypothesize that such height potential is related to tolerance mechanisms. Parker and Gilbert (2018) also reported that the impact of infection (tolerance) on 17 closely related clover species was less negative on fast-growing species, possibly because of their better ability to acquire resources in the environment and compensate for damage (de Jong 1995). Moreover, some authors suggested that tolerance could be especially advantageous for long-lived species (Roy et al 2000). Although tolerance has been far less investigated than resistance, there is ample evidence of its occurrence in crops and wild plants (Pagan & Garcia-Arenal 2020), and more in-depth molecular approaches would probably be needed for unraveling the cascades of metabolic pathways behind tolerance and its correlation with growth-related traits (Monson et al 2021; Monson et al 2022).

### 3. Increased powdery mildew pressure had no equalizing effect on the relative contribution of mother trees to the next generation

The surviving population was slightly less diverse in terms of family composition under high than under low powdery mildew pressure. This pattern may be explained by previous results showing that the advantage of the fast-growing families over the slow-growing families in terms of survival was not suppressed under pathogen exposure. On the contrary, fast-growth might be associated with higher tolerance to infection damage. Parker and Gilbert (2018) obtained very similar patterns with closely related species instead of families and suggested that greater tolerance in fast-growing species may limit rather than promote species coexistence. Similarly, at infra-specific level, Mundt et al (2008) showed that the absolute fitness advantage of the more competitive genotype in absence of disease increased in the presence of disease.

However, differences in mean final height among families were reduced under disease pressure, as well as the variance within families. During the time frame of our experiment, this did not affect the family ranking for survival between disease exposures but maybe in a longer term, or with greater disease pressure, powdery mildew could have an equalizing effect on family and individual performances (survival).

### 4. Impact of powdery mildew on genetic diversity

We did not observe any changes in genetic diversity (estimated by the mean H_e_ across a large number of SNPs) or any significant genetic differentiation (estimated by F_ST_) between the initial and the surviving oak populations in both exposures, with mortality rates as high as 60% in some naturally infected plots at the end of the monitoring period. Few studies have addressed the impact of disease on genetic diversity of natural plant populations contrary to animal populations (McKnight et al 2017). One of the best studied wild plant pathosystem in a long time series is the interaction between *Linum marginale* and *Melampsora lini*, which showed temporal patterns of genetic change in the host and pathogen at local scale, consistent with coevolutionary dynamics (Thrall et al 2012). However, these changes were associated with a gene-for-gene model, *i.e.,* the existence of matching genes for resistance in the host and virulence in the pathogen (Flor 1971), which is not characterized for the oak-powdery mildew interaction. One possible explanation to the lack of any observed change in our study is that the powdery mildew pressure was not strong enough to have significantly affected the very high background diversity revealed across the oak genome (Plomion et al 2018). Also, few if any powdery mildew infection causative or linked variants are probably included in our SNP sets due to a very low background linkage disequilibrium across the genome (Lang et al 2021). A similar argument can be invoked for variants involved in the genetic determinism of traits linked to growth and survival that are most probably multigenic, which means that one single allelic variant explains a very small part of the total variation (see below in section 5).

We also did not observe a significant increase of the mean individual heterozygosity in surviving populations compared to the initial ones, or between surviving populations between exposures. An increase of this index could have been linked to an overall Heterozygosity-fitness correlation (HFC), associated with a deleterious effect of inbreeding (Slate et al 2004). We can notice, however, that the seedlings with lowest heterozygosity values (PHt inferior to 0.235) were often dead at the end of the experiment, leading to a slight increase of this statistic at the end of the experiment for most families in both exposures. Although a slight effect of multilocus heterozygosity at SSRs was detected on growth traits in *Q. robur* (Vranckx et al 2014b, less than 5% of total variation explained), and in other species with different reproductive strategies (Cole et al 2016 in Aspen; Stilwell et al 2003 in *Castanea dentata*), a better resistance or tolerance to pathogen infection for heterozygotes has not been established for plants, contrary to animals (*e.g.*, Budischak et al 2023). Specifically, oaks have generally large populations with a low inbreeding of their seedlings (Gerzabek et al 2020), a genetic context for which HFC is not expected (Slate et al 2004). HFC may also be more easily detected in stressful environments (Mopper et al 1991, and see references above). In addition, opposite correlations between heterozygosity, growth, and mechanisms of resistance against pathogens were reported in Cole et al (2016). Antagonistic effects of heterozygosity on different biological traits could occur in oak seedlings, thus masking possible HFC. As explained above (section 2), tolerance to oak powdery mildew could be a composite process involving different physiological functions that would lead to moderate optimizing selection effects on genotypes and thus the maintenance of a mean diversity level (Walsh & Lynch 2018).

### 5. Few genetic associations identified

Our design with well-characterized phenotypic traits and individuals genotyped at several hundred SNPs was suitable to test for genetic associations. Overall, in conservative statistical testing conditions, very few significant associations were observed: zero, one or two per trait for survival, mean powdery mildew infection and height, and up to seven for acorn weight, across both exposures. We did detect one SNP significantly associated with seedling survival in the natural exposure, which differed from the one detected in the disease protected exposure. This SNP was not associated with powdery mildew infection, and the allele with a beneficial effect on survival was at very low frequency. If true, this beneficial effect might be counterbalanced by pleiotropic effects, and the advantage detected in our study might be related to particular conditions. Indeed, this SNP was located in a gene coding for a putative histone H4, a type of protein involved in the structure of chromatin that has been linked to survival strategy against drought in plants (Kim et al 2017). The second SNP associated with survival but under protected exposure was located in a gene coding for a family of proteins involved in organelle biogenesis (O’Toole et al 2008).

Genetic association studies of resistance to plant diseases are common in pathosystems of agronomic interest (*e.g.*, maize (Zhao et al 2022, reviewed in Shrestha et al 2019), wheat (Du et al 2021) or soya bean (Wen et al 2018)). They are far less common in natural pathosystems. The genetic architecture of resistance (*sensu lato*) to powdery mildew in oak species is still poorly understood, although QTLs have been detected and some candidate genes have been suggested (Bartholomé et al 2020). The present study points to another possible candidate gene identified by the GWAS approach, a gene coding for a putative ethylene response factor C3. Interestingly, a similar type of protein has been linked to pathogen resistance in cotton (Guo et al 2016). Despite a measurable effect on tree mortality from oak powdery mildew in our experimental setup, the only SNP associated with susceptibility to powdery mildew was not associated with the survival trait. This is probably related to the complex and partly indirect nature of the effect of powdery mildew on oak mortality, as evidenced by the SEM model results.

## Acknowledgements

We are very grateful to Rémy Petit and Christophe Plomion for insightful discussions at the initiation of this study. We sincerely thank Inge van Halder and several technicians and students who helped in the set up and monitoring of the field design, with a special contribution of the Unité expérimentale Forêt Pierroton, UE 0570, INRAE, especially Frédéric Bernier, Luc Puzos and Henri Bignalet. We thank Marie-Christine Le Paslier, Dominique Brunel and the Genoscope for advice and for performing the SNP genotyping.

## Data, scripts and codes availability

Data, supplementary data and scripts are available online in a Zenodo repository: https://doi.org/10.5281/zenodo.7517641

## Supplementary material

Supplementary Materials are available online: https://doi.org/10.5281/zenodo.7931510

